# DiMeLo-cito: a one-tube protocol for mapping protein-DNA interactions reveals CTCF bookmarking in mitosis

**DOI:** 10.1101/2025.03.11.642717

**Authors:** Nathan Gamarra, Cy Chittenden, Kousik Sundararajan, Jacob P. Schwartz, Sofia Lundqvist, Denise Robles, Oberon Dixon-Luinenburg, Jeremy Marcus, Annie Maslan, J. Matthew Franklin, Aaron Streets, Aaron F. Straight, Nicolas Altemose

## Abstract

Genome regulation relies on complex and dynamic interactions between DNA and proteins. Recently, powerful methods have emerged that leverage third-generation sequencing to map protein-DNA interactions genome-wide. For example, Directed Methylation with Long-read sequencing (DiMeLo-seq) enables mapping of protein-DNA interactions along long, single chromatin fibers, including in highly repetitive genomic regions. However, DiMeLo-seq involves lossy centrifugation-based wash steps that limit its applicability to many sample types. To address this, we developed DiMeLo-cito, a single-tube, wash-free protocol that maximizes the yield and quality of genomic DNA obtained for long-read sequencing. This protocol enables the interrogation of genome-wide protein binding with as few as 100,000 cells and without the requirement of a nuclear envelope, enabling confident measurement of protein-DNA interactions during mitosis. Using this protocol, we detected strong binding of CTCF to mitotic chromosomes in diploid human cells, in contrast with earlier studies in karyotypically unstable cancer cell lines, suggesting that CTCF “bookmarks” specific sites critical for maintaining genome architecture across cell divisions. By expanding the capabilities of DiMeLo-seq to a broader range of sample types, DiMeLo-cito can provide new insights into genome regulation and organization.

## Introduction

Chromosomes store genetic information encoding a diverse set of cellular programs to ensure proper organism development, survival, and reproduction. An outstanding goal of biology is to fully understand how this information is accessed and regulated at the molecular scale and across cellular generations. In the cell, DNA is regulated in large part through the highly ordered assembly of protein complexes at precise locations in the genome. Many techniques have been developed to assess genome-wide binding of target proteins using short-read DNA sequencing, but they have important limitations, such as the inability to map within repetitive regions. ChIP-seq, perhaps the most widely used method ^1^, relies on chemical fixation of molecules to DNA followed by DNA fragmentation, antibody-based enrichment, crosslink reversal, and library preparation for DNA sequencing. Fixation is known to introduce substantial artifacts, most notably during mitosis ^2^, which can obscure the true distribution of DNA binding. Native ChIP-seq does not require fixation but is only effective with very large cell inputs and very high-affinity targets such as histones^3^. More recent methods, such as short-read Chec-seq ^4^ and CUT&RUN ^5^, instead retain condensed chromatin within permeabilized nuclei ^6^. Native chromatin can then be incubated with antibodies, which are targeted by Protein A-linked nucleases that release short DNA fragments, which are then amplified and sequenced. However, between incubations, excess unbound antibody/enzyme complexes must be washed away and buffers must be exchanged, which is achieved by centrifugation and aspiration. Although versions of such nuclease protocols have been attempted on mitotic cells with variable success ^7,8^, it is unclear how efficiently chromosomes are retained without the nuclear envelope and lamina, which are disassembled during mitosis ^9^.

We recently developed Directed Methylation with Long-read sequencing (DiMeLo-seq), which enables the interrogation of protein-DNA interactions across the genome using third-generation sequencing reads ranging from 2 kb to 25 kb in the case of Pacific Biosciences HiFi sequencing to as large as 1 Mb with Oxford Nanopore Technologies (ONT) ultralong sequencing ^10^. Like CUT&RUN, DiMeLo-seq utilizes antibodies to bind target molecules within permeabilized nuclei, which are subsequently targeted by Protein A (pA) fused to an enzyme. For CUT&RUN and similar protocols, this pA-fused enzyme is a nuclease, which fragments nearby DNA into short pieces. For DiMeLo-seq and similar protocols^11,12^, this pA-fused enzyme is a DNA adenine methyltransferase derived from bacteria, which is activated upon addition of the methyl donor S-Adenosyl Methionine (SAM). Since 6mA is virtually absent from vertebrate DNA, the positions of protein binding sites can be directly and quantitatively inferred by the frequency of 6mA modification, without requiring DNA fragmentation. DiMeLo-seq was recently employed to map the localization of CTCF, CENP-A, H3K9me3, H3K27me3, H3K27ac, H3K4me3, FOXA1, FOXA2, TEAD1 and LMNB1 genome-wide ^10,13–17^. Third-generation sequencing of native DNA molecules also unlocks information previously inaccessible to short-read methods, such as the relationship of binding events with respect to endogenous 5mCpG modifications. ONT nanopore sequencing enables the generation of ultralong (>150 kb) DNA sequencing libraries, which can be mapped to large, highly repetitive regions of the genome and to parental haplotypes. Through the use of DiMeLo-seq coupled with ONT sequencing, we recently profiled CENP-A distributions within human centromeres and confirmed that CENP-A assembles on hypomethylated regions within alpha-satellite higher-order repeat arrays^10^.

While DiMeLo-seq represents a significant advance, it does have some important limitations. Because DiMeLo-seq relies on sequencing native, unamplified DNA, the yields of DNA following purification are critical to the success of the method. DiMeLo-seq, like CUT&RUN, also relies on the nuclear lamina to enclose the genome for pelleting and wash steps, which are lossy and contribute to the reduction in DNA recovery at the end of the protocol. Consequently, an approach without wash steps would likely improve DNA yields. Furthermore, reducing or eliminating the number of wash steps reduces the possibility for sample damage and loss of protein binding. Challenging a wash-free approach is the fact that the simple addition of antibody and Protein A to the reaction at concentrations required for efficient target binding would result in the aggregation and precipitation of antibody-pA complexes. This is because pA is a multivalent binder of antibodies ^18^, which, coupled with the hetero-tetrameric structure typical of antibodies, enables aggregation and precipitation. As a result, removing wash steps would require a substantial redesign of the core DiMeLo-seq protocol.

Here, we present DiMeLo contained in tube one (DiMeLo-cito), a streamlined protocol that enables the mapping of arbitrary targets in the genome in a single tube without wash steps. This is accomplished by a careful redesign of the original DiMeLo-seq protocol including optimization of the biochemical activity of its DNA adenine methyltransferase, Hia5^19^. We demonstrate the utility of DiMeLo-cito by mapping the genome-wide binding of the protein CTCF during interphase and mitosis. DiMeLo-cito substantially expands the efficiency and utility of DiMeLo-seq, requiring less hands-on time, less starting material, and fewer constraints on input sample types. We envision DiMeLo-cito replacing DiMeLo-seq for most applications.

## Results

### Development of an optimized single-tube protocol for DiMeLo-seq

The standard DiMeLo-seq protocol involves 4 major steps prior to DNA extraction and sequencing: 1) nuclear permeabilization, 2) primary antibody incubation, 3) pA-Hia5 incubation, 4) SAM incubation to activate methylation ^10,13^. Washes or buffer exchanges are required between each of these steps to maintain optimal conditions for each incubation and to remove excess antibodies and enzyme. In order to develop DiMeLo-cito, we had to devise 3 key innovations. First, we conceived of a protocol that would retain optimal buffer conditions by successively adding and diluting buffer components rather than performing buffer exchanges (Figure 1A)^20^. Second, to prevent aggregation of antibody complexes, we replaced Protein A fusions of Hia5 with recently-developed nanobodies that bind antibodies with species specificity ^21^. Since nanobodies are monovalent binders of their antigen, bridging between antibodies is prevented even at very high concentrations of both components ^22^. Finally, we devised a simple centrifugation-free approach to replace washes and deplete excess antibody and nanobody-Hia5 (Nb-Hia5). We did this by enclosing an affinity resin in a semipermeable membrane that permits the passage of the relatively small antibody-Nb-Hia5 complexes but prevents the direct interaction of chromatin with the affinity resin (Figure 1A). We call this membrane-enclosed affinity resin device an Enclosure of Micron-scale Particles with No Adhesion of DNA Analyte (EMPaNADA). By incubating the reaction with an EMPaNADA, we effectively deplete excess antibody-Hia5 complexes that are not bound to chromatin, while ensuring maximal recovery of DNA for downstream sequencing.

**Figure 1.**
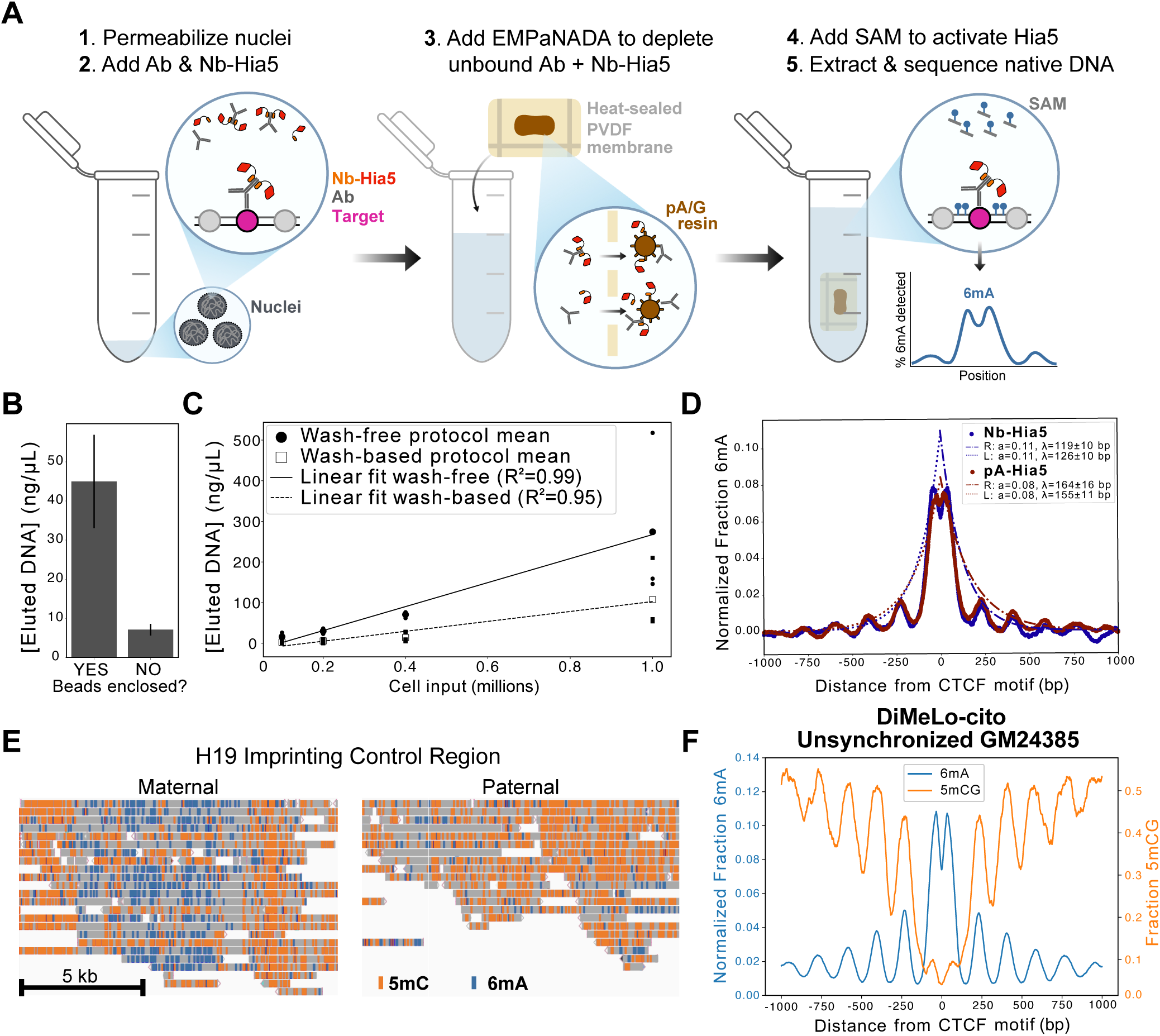
Genome-wide mapping with enhanced DNA recovery using DiMeLo-cito. **A.** Schematic overview of the DiMeLo-cito workflow. Permeabilized nuclei are incubated with antibodies that bind target molecules in the genome, which in turn are bound by a nanobody-fused Hia5 adenine methyltransferase. Instead of washing, the sample is diluted and incubated with an affinity resin enclosed in a semipermeable membrane in an assembly known as an EMPaNADA to remove free enzyme-antibody complexes. After activation of Hia5 with SAM, nearby adenines are methylated, and DNA is directly extracted and subjected to long-read library preparation and sequencing. **B.** DNA extraction recoveries from ∼1 million digitonin-permeabilized nuclei in the presence of paramagnetic affinity resins directly added to the reaction or enclosed in an EMPaNADA. Samples were eluted from beads with 100 µL buffer. **C.** DNA extraction recoveries from varying inputs of permeabilized nuclei subjected to the washing protocol of DiMeLo-seq or diluted with an EMPaNADA and kept in a single tube. **D.** Methyladenine enrichment profiles at bound CTCF sites for DiMeLo-seq and DiMeLo-cito targeting CTCF and sequenced to ∼1X coverage genome-wide. Plots were generated with a 50 bp smoothing window. Peaks in methyladenine signal were fit to exponential decays. Fit parameters for the maximal value (a) and the decay constant (lambda) are shown in the figure legends. **E.** Screenshot from IGV at the H19 imprinting control region with reads aligned to the HG002v1.1 genome assembly. Locations of methyladenine and methylcytosine basecalls with a confidence threshold >0.99 are shown. The regions shown correspond to chr11_MATERNAL:2,065,542-2,078,512 and chr11_PATERNAL:2,057,830-2,069,830. **F** Methyladenine and methylcytosine distributions at bound CTCF sites with DiMeLo-cito sequenced to approximately 30X coverage. Methyladenine signal was subtracted from nontargeting (IgG isotype) controls.

### Optimization of reaction buffer

We calculated buffer volumes to enable all steps of the protocol to take place within the constraints of a 2 mL microcentrifuge tube (Figure 1A), while adding and diluting salt and detergent to maintain optimal conditions for each incubation step without buffer exchange. To begin buffer optimization, we purified two nanobody-Hia5 (Nb-Hia5) fusion constructs, each targeting antibodies produced in rabbit and mouse ^21^ (Figure 1 S1A). Success in purification relied on the presence of an N-terminal fusion of Maltose Binding Protein (MBP), which improved the solubility of Nb-Hia5 fusions. We then tested these preparations in their ability to methylate a DNA probe in vitro using a restriction enzyme protection assay. We found that both Nb-Hia5 constructs were highly active in their methyltransferase activity and that this was reproducible between enzyme preparations (Figure 1 S1B).

In order to achieve the most robust enzyme activity, we considered buffer components that might promote Hia5’s methyltransferase activity. Use of organic anions in buffers more accurately mimics the intracellular environment, whose counterion composition is primarily organic acids and phosphates ^23^. Furthermore, organic anions have been shown to enhance DNA binding of mammalian and prokaryotic proteins in vitro ^24^. We found that replacing the standard chloride-based buffers used in DiMeLo-seq with acetate or glutamate improved the activity of Hia5 (Figure 1 S1C). DNA adenine methyltransferases have previously been shown to be sensitive to the reducing potential of the buffer used ^25^. We found that the inclusion of the reducing agent DTT at an optimal concentration of ∼1-2 mM increased Hia5 activity ∼2-fold (Figure 1 S1D). Initially we were concerned that reducing agents would disassemble antibodies due to their cleavage of disulfides that stabilize antibody structure ^26^. However, size exclusion chromatography analysis suggested that the antibodies used in this study retain their structure in spite of disulfide bond scission even at concentrations as high as 25 mM (Figure 1 S1E). These data suggest that noncovalent interactions between the heavy and light chains of antibodies are sufficient to retain antibody quaternary structure.

### Replacement of washes with affinity depletion

With an optimized buffer system in place, we explored strategies to remove free enzyme complexes from solution without requiring washes. Affinity resins, which selectively bind and precipitate target proteins, are widely used for depleting enzymes from complex mixtures ^27^. We first tested paramagnetic beads targeting the MBP tag on Nb-Hia5 (anti-MBP) using an in vitro restriction enzyme cleavage assay to measure their ability to deplete methyltransferase activity. Indeed, free-floating beads mixing in solution rapidly and effectively depleted all detectable methyltransferase activity (Figure 1-S2A). However, when beads were incubated freely and directly with nuclei, followed by magnetic separation prior to nucleic acid extraction, DNA extraction yields were poor, likely due to the adhesion of nuclei to paramagnetic beads (Figure 1B). To mitigate harmful interactions with nuclei, we explored retaining beads against the tube wall using a magnet throughout the bead incubation. However, this approach resulted in poor depletion efficiency, as determined by a restriction activation assay, likely due to the occlusion of the surface area for bead binding.

Seeking to protect nuclei and DNA from bead binding while maximizing binding capacity, we devised an alternative strategy: enclosing the beads within a semipermeable membrane that allows enzyme complexes to diffuse through while excluding nuclei. We call this assembly an EMPaNADA. To make EMPaNADAs, we folded beads into a 0.2-micron pore-diameter polyvinylidene fluoride (PVDF) membrane, which was then heat-sealed to prevent bead leakage (see Methods). The selected pore size permits the diffusion of antibody-Nb-Hia5 complexes (∼1–100 nm in diameter) across the membrane, while restricting nuclei (∼2–10 microns in diameter). Incubation of nuclei with an EMPaNADA effectively depleted methyltransferase activity over ∼4 hours at room temperature (Figure 1 S2 B-D). Furthermore, we found that incubating permeabilized nuclei with an EMPaNADA substantially increased DNA recovery compared to free-floating beads, with an ∼80% yield (Figure 1B).

To accelerate enzyme depletion, we considered alternative affinity resins that may more rapidly deplete Hia5-antibody complexes. We considered whether antibodies might hinder the efficiency of anti-MBP beads in removing enzyme-antibody complexes. As an alternative, we tested Protein A/G (pA/G)-linked resin, which binds antibodies with picomolar affinity and has also been reported to interact with some nanobodies ^28^. We compared the depletion kinetics of pA/G EMPaNADAs with anti-MBP EMPaNADAs and found that Nb-Hia5 is depleted approximately tenfold faster with pA/G resin (Figure 1 S2C-D). We also found that pA/G resin directly binds both nanobodies used in our Nb-Hia5 constructs (Figure 1 S2E). Given these results, we investigated whether combining anti-MBP and pA/G assemblies could further improve depletion. Surprisingly, while the kinetics remained similar to pA/G EMPaNADAs alone, the equilibrium concentration of soluble methyltransferase was higher, resulting in higher background activity (Figure 1 S2C-D). We speculate that the inclusion of anti-MBP in addition to pA/G resin increases the equilibrium concentration of nanobody-Hia5 complexes in solution, resulting in higher background activity.

### Testing an initial protocol

Combining all of the insights above, we generated a final protocol for DiMeLo-cito (Figure 1A and Methods). First, isolated cell pellets are treated with the saponin detergent digitonin to permeabilize cell and nuclear membranes. Permeabilized nuclei are then incubated with primary antibody and Nb-Hia5 to bind targets in the genome. The sample is then diluted 10-fold in activation buffer (minus SAM), and incubated for 2 hours at room temperature in the presence of an EMPaNADA. Following depletion, methylation is initiated by addition of SAM and DTT for 2 hours at 37°C.

To benchmark the performance of DiMeLo-cito, we performed the protocol with ∼1 million GM24385 transformed lymphoblastoid cells (LCs) using a CTCF targeting (anti-CTCF) or nontargeting antibody (IgG). We then performed nanopore sequencing and analyzed the degree of specific methylation detected at the highest-occupied ChIP-peaks derived from ENCODE ChIP-seq datasets of GM12878 LCs ^29^. First we sought to determine the consequence of performing non-targeted methylation using an IgG isotype control antibody without depletion of excess enzyme. As expected, substantial signal was observed at CTCF sites as they are accessible to modification by free Hia5 enzyme ^30^ (Figure 1 S2F). We then tested the impact of free antibody-enzyme depletion in the presence of freely diffusing beads or beads enclosed in an EMPaNADA. We found that incubation of free beads or an EMPaNADA resulted in depletion of non-targeted methyltransferase activity at CTCF sites (Figure 1 S2F). Interestingly, incubation of free Nb-Hia5 without IgG resulted in reduced depletion efficiency (Figure 1 S2F). We also found that using two EMPaNADAS containing pAG and MBP produced more nontargeted methylation at CTCF sites (Figure 1 S2G), consistent with the reduced depletion observed in restriction digestion assays.

### Properties of DiMeLo-cito

After observing that EMPaNADAs successfully reduce non-targeted DNA methylation, we sought to determine whether targeted CTCF signal could be detected using DiMeLo-cito. Using either anti-MBP or pA/G resin, we observed substantial (≥2-fold) increase in methyladenine frequency with anti-CTCF compared to IgG (Figure 1 S3A). Consistent with restriction assays, we found that anti-MBP resin results in higher non-specific methylation when compared to pA/G resin (Figure 1 S3A). To verify that marking by Nb-Hia5 is specific to its nanobody targeting domain, we compared the frequency of targeting with anti-mouse or anti-rabbit nanobody-Hia5 using anti-CTCF antibodies produced in cognate or noncognate species. Indeed, we observed that marking of CTCF sites was more enriched when nanobody-Hia5 was incubated with its cognate antibody as opposed to its noncognate antibody (Figure 1 S3B). This suggests that nanobody-Hia5 engages its target antibody with species specificity.

Having observed that DiMeLo-cito successfully enriches methyladenine at targeted sites, we then sought to determine the features influencing antibody-targeted signal. We first wondered if the degree of specific signal was sensitive to the duration of methyl transfer. To test this, we varied the duration of incubation with SAM at 37°C from 30 minutes to 3 hours and compared the signal at CTCF sites with anti-CTCF and IgG. We found that the frequency of adenine methylation at CTCF sites increased monotonically with time in both anti-CTCF and IgG conditions, leveling off at approximately 2 hours (Figure 1-S3C). However, the ratio of targeting and non-targeting methylation remained roughly constant (∼3-4-fold) regardless of the duration of incubation (Figure 1 S3C).

Methylation of DNA by Hia5 is strongly sensitive to the packaging state of DNA into chromatin^30^. CTCF sites generally are accessible to marking by methyltransferases, explaining the local increase in methylation in non-targeting conditions. It is therefore possible that the optimal marking duration may be dependent on the targets marked. To determine how DiMeLo-cito performs at specifically marking DNA hyper-accessible regions of the genome, we performed the protocol targeting anti-rabbit nanobody-Hia5 to the histone modification H3K4me3, which resides at active transcription start sites (TSSs) ^31^. We then compared the frequency of methyladenine at the top quartile of TSSs with anti-H3K4me3 and IgG across varying incubation times with SAM. As with the CTCF targeting case, we found that overall methylation increased with time, while the ratio of targeted and non-targeted methylation remained roughly constant at ∼2-fold (Figure 1 S3D). Together, these data suggest that the degree of methylation is dependent on both the incubation time and the loci targeted. However, for a particular set of loci, the fraction of methylation that is due to targeting is approximately constant over a 3 hour incubation period.

To directly compare the properties of Nb-Hia5 and pA-Hia5 in marking the genome, we compared datasets at ∼1X coverage with Nb-Hia5 using the DiMeLo-cito protocol with pA/G beads and the standard DiMeLo-seq protocol using pA-Hia5 in the same activation buffer as Nb-Hia5. At higher depth, DNA methylation by Hia5 flanking CTCF sites becomes more apparent and produces a regular oscillating pattern ^10^. This pattern is due to the regular positioning of nucleosomes flanking CTCF sites, where the valleys of methylation correspond to a positioned nucleosome blocking underlying DNA methylation ^32^. Additionally, a valley at the center of the CTCF motif is apparent due to the footprinting of its site due to binding. Fitting the peaks in methylation that regularly flank the CTCF binding site to an exponential decay and comparing fit parameters permits the quantitative comparison of both the peak methylation at the bound site of the tethered enzyme complex and its degree of spatial constraint. Doing this analysis suggests that Nb-Hia5 modifies chromatin flanking CTCF sites ∼40% more efficiently (determined by peak methylation) and with a ∼25% tighter distribution (determined by decay constants) (Figure 1 C). These data suggest that DiMeLo-cito using Nb-Hia5 specifically marks CTCF sites with greater spatial constraint and slight increases in enzyme efficiency.

### Improvements in DNA extraction

By constraining our protocol to a single tube without washes, we anticipated an improvement of the maximal recovery of input DNA. To test this possibility, we varied the cell input and subjected it to permeabilization and washing according to the standard DiMeLo-seq protocol, or to dilution, EMPaNADA incubation, and extraction according to the DiMeLo-cito protocol. We observed an ∼2-fold increase in DNA recovery across dilutions, indicating that the wash-free DiMeLo-cito protocol loses fewer nuclei. Given this, we sought to determine if fewer than 1 million cells could be used as input for DiMeLo-cito. From an input of 100,000 GM24385 cells, we were able to generate low-coverage (∼0.1X) ONT sequencing data using either anti-CTCF or IgG control antibody. We observed methyladenine enrichment at highly bound CTCF loci with targeting antibody but not with control antibody (Figure 1 S2E). We expect that future improvements to ONT’s library prep and sequencing efficiency will reduce these input requirements further.

A major feature of ONT sequencing is the ability to sequence ultralong DNA fragments with median fragment lengths in excess of 100 kb ^33^. In the process of developing DiMeLo-cito, we sought approaches that would improve the speed and simplicity of extracting ultra high molecular weight (UHMW) DNA fragments. Typical protocols for UHMW DNA purification rely on the precipitation of large DNA fragments onto silica, gentle washing of the precipitated DNA to remove contaminants, and finally resolubilization in buffer. Resolubilization is frequently rate limiting due to the slow rehydration kinetics of precipitated DNA, often taking days to fully solubilize. Previous studies suggest that small zwitterions, molecules with positively and negatively charged moieties but no net overall charge, improve the solubilization of DNA into buffer by altering the dielectric constant of water ^34^. We were curious whether 6-amino caproic acid (6-ACA), a zwitterion structurally similar to the side-chain of lysine, could speed the solubilization of UHMW DNA fragments. We found that including 0.5M 6-ACA during solubilization dramatically accelerated solubilization of ethanol-precipitated UHMW DNA (Figure 1 S4A).

We then sought to determine whether DiMeLo-cito is compatible with extraction of UHMW DNA in the presence or absence of 6-ACA. We subjected DiMeLo-cito samples to UHMW DNA extraction using either a modified commercial kit (NEB UHMW DNA extraction kit) or an in-house DNA extraction method (see Methods), which used 6-ACA to solubilize extracted DNA. We compared the resulting fragment length distributions of the purified DNA using the Agilent Femto Pulse capillary electrophoresis system (Figure 1 S4B). We found that both methods extracted UHMW DNA with median fragment lengths of ∼400 kb (Figure 1 S3B-C). This suggests that DiMeLo-cito generates DNA of a quality suitable for ultralong DNA libraries. Furthermore, it also suggests that 6-ACA does not negatively impact fragment lengths of recovered DNA. Importantly, we also found that DNA extracted with 6-ACA generates high-quality ONT sequencing data (Figure 1 S4D-E). These data suggest that DiMeLo-cito is compatible with the extraction of UHMW DNA and that the use of 6-ACA to improve solubilization is compatible with ultralong ONT library preparation and ONT sequencing.

### Validation of genome-wide mapping with haplotype phasing

Confident that DiMeLo-cito recapitulates previous observations of CTCF binding, we generated a deeply sequenced ONT dataset at approximately 30X coverage in 1 million unfixed GM24385 cells. The datasets were of high quality with a modal Phred Q score of ∼21 and a median fragment length of ∼14 kb (Figure 1S4D-E). As previously observed, we also see several features associated with CTCF binding including the oscillating levels of methylation proximal to the CTCF site due to nucleosome phasing at bound sites, a precipitous drop in CpG methylation approaching the CTCF bound site, and a focal drop of 6mA signal over the motif, which is footprinted by CTCF binding ^10^. With this deep dataset, we found that there is >2-fold enrichment of targeted DNA methylation compared to non-targeting datasets. This ratio was largely insensitive to the choice for adenine methylation confidence threshold over a range of 0.75-0.98 (Figure 1 S4F). To evaluate whether DiMeLo-cito recapitulates the quality of mapping to specific haplotypes, we mapped reads to the haplotype-phased HG002 genome assembly generated from GM24385 LCs. At the H19 imprinting control region on chromosome 11, CTCF specifically binds only to the maternal allele due to paternal-specific CpG methylation that blocks CTCF binding. Binding of CTCF to the maternal allele insulates a distal enhancer that blocks expression of the gene Igf2. Consistent with several studies as well as previous observations with DiMeLo-seq ^10^, we observe substantial methyladenine enrichment on the maternal allele with substantial depletion of CpG methylation (Figure 1D). In contrast, on the paternal ICR, we observe substantial CpG methylation but highly depleted CTCF binding (Figure 1D). Together these data suggest that DiMeLo-cito maps CTCF binding to phased haplotypes and recapitulates existing distributions.

### DiMeLo-cito reveals mitotic CTCF bookmarking in non-cancer-derived cell types

With an optimized protocol for DiMeLo-cito, we sought to apply it to study mitotic chromosome architecture. To generate a highly purified population of mitotic GM24385 LCs, we employed a double thymidine block procedure followed by a nocodazole arrest in early mitosis, before washout and harvesting in HBSS containing MG132 to prevent progression through metaphase. This generated a highly purified population (∼90% mitotic by flow cytometry, Figure 2 S1A) of mitotic LCLs in prometaphase (Figure 2 S2A). To further verify mitotic enrichment, we performed DiMeLo-cito in unsynchronized and mitotic cells and compared the chromosomal distribution of H3S10ph, a strong marker of cells actively undergoing mitosis ^35^. Recent studies suggest that in interphase H3S10ph is found at promoters and during mitosis it increases and spreads into gene-rich locations to promote chromosome compaction^36^. By DiMeLo-cito we detected enrichment of H3S10ph near active transcription start sites and gene bodies, which increases in the mitotic cell sample relative to the unsynchronized sample (Figure 2 S1B-C). This suggests that DiMeLo-cito can recapitulate known patterns of mitotic chromatin binding with our mitotic cell sample.

**Figure 2.**
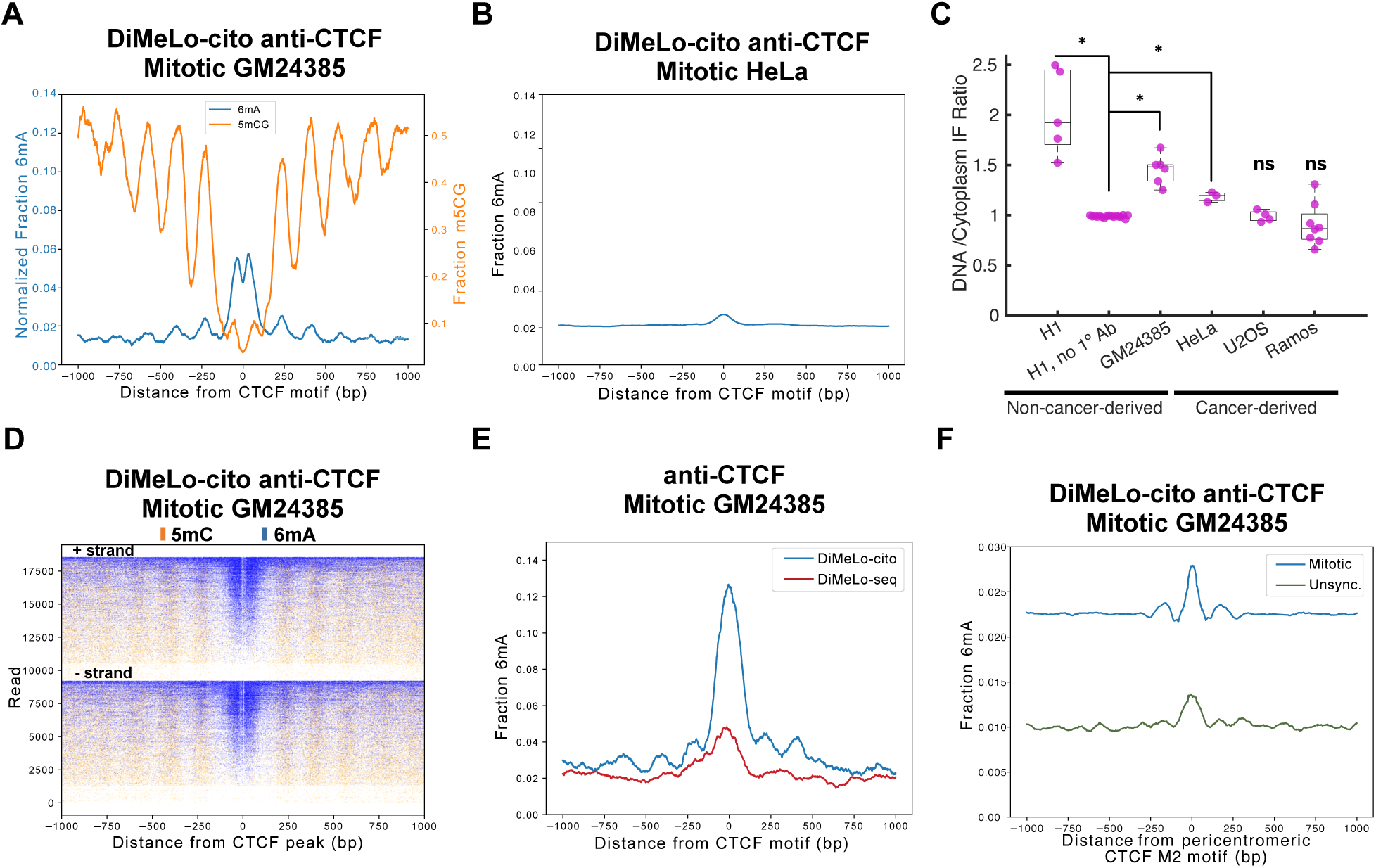
DiMeLo-cito reveals CTCF binding in unfixed mitotic GM24385 LCs. **A.** Methyladenine and methylcytosine distributions from DiMeLo-cito targeting CTCF in unfixed mitotic GM24385 with nontargeting background subtracted. DNA from 1 million cells was sequenced to ∼30X coverage and plots were generated with a 30 bp smoothing window **B.** Methyladenine distribution from DiMeLo-cito targeting CTCF in unfixed HeLa cells synchronized in mitosis sequenced to ∼2X coverage. Plots were generated with a 50 bp smoothing window. **C.** Quantification of immunofluorescence (IF) signal targeting CTCF in various unfixed cell lines. Asterisks denote p-values <10^-10^ for a two-sample tailed t-test calculated between the secondary-only control condition and each cell line. **D.** Plots of single reads from B centered on CTCF sites and separated based on strand and sorted in descending order by the fraction of the read containing methyladenine. **E.** Comparison of standard DiMeLo-seq targeting CTCF in mitotic cells at ∼0.1X coverage and DiMeLo-cito data in B downsampled to 0.1X coverage for comparison. Both plots were smoothed with a 50 bp smoothing window. **F.** An enrichment profile plot of CTCF signal at M2 motifs within centromeric transition zones during mitosis (blue) or in unsynchronized cells (green).

Previous studies have indicated that nearly all CTCF is shed from mitotic chromosomes in human cells in prometaphase^7^, with the potential exception of specialized pericentromeric sites^37^. However, these studies were done in cancer cell lines, which may not be representative of non-cancerous human cells. Further, studies of mouse ESCs suggest that substantial amounts of CTCF associate with chromosomes throughout mitosis^38^. To determine the extent of mitotic chromosome binding by CTCF, we performed DiMeLo-cito targeting GM24385 LCs synchronized in mitosis and sequenced to ∼10X coverage. We observed strong enrichment of 6mA at CTCF sites that are highly bound by CTCF in unsynchronized cells (Figure 2A), although the peak normalized modification frequency was reduced approximately ∼50% compared to unsynchronized cells (Figures 1F,2A). We also observed several features of CTCF-bound sites including phased nucleosomes and an anticorrelation with CpG methylation (Figure 2A,D).

Importantly, we were unable to detect mitotic CTCF binding in the same sample performing standard DiMeLo-seq (Figure 2E). We speculate that the wash-free protocol of DiMeLo-cito better preserves CTCF binding to unfixed mitotic chromosomes, explaining its detection only with DiMeLo-cito. To determine if DiMeLo-cito recapitulates previous studies showing low CTCF enrichment in mitosis in other cell types ^7^, we performed DiMeLo-cito targeting CTCF on HeLa cells synchronized in mitosis. This resulted in only faint 6mA enrichment across CTCF ChIP-seq peaks from the same cell type ^29^ (Figure 2B). Together, these results suggest that CTCF binding in mitosis is highly depleted at major interphase-bound sites in HeLa cells but only modestly reduced at bound sites in GM24385 LCs.

To further test whether CTCF was bound in our mitotic cell samples, we performed immunofluorescence (IF) staining on unfixed mitotic GM24385 cells with confocal imaging, and we observed abundant CTCF staining across the entire chromosome (Figure 2 S2A). We also decided to look for CTCF binding using IF in pluripotent H1 human Embryonic Stem Cells (hESCs). Using very gentle permeabilization and incubation conditions without fixation or synchronization, we observed robust staining of CTCF across H1 mitotic chromosomes (Figure 2 S2A-B). This suggests that CTCF binding during mitosis is not limited to EBV-immortalized LCs and is a feature of cultured pluripotent cells.

To compare mitotic chromosome binding of CTCF in previously studied cancer cell lines, we synchronized U2OS and HeLa cells in mitosis and subjected them to IF staining for CTCF without fixation. Consistent with previous studies of mitotic chromosomes in these lines ^7^, we observed essentially no CTCF staining for U2OS chromosomes and limited staining for HeLa chromosomes (Figure 2 S2A-B). To determine if CTCF binding is lost in cancerous cells of similar origin as GM24385 LCs, we performed IF staining in unfixed Ramos cells, which were derived from a Diffuse Large B-Cell Lymphoma sample. Strikingly, we observed essentially no staining of CTCF on the mitotic chromosomes of Ramos cells, suggesting that B-cells of different states can harbor different mitotic CTCF distributions (Figure 2 S2A-B). Individual confocal slices from multiple mitotic cells from all of the samples described were subjected to quantification, revealing statistically significant differences in staining intensity to mitotic chromosomes (Figure 2C). Together, these data suggest that CTCF binding to human mitotic chromosomes is more widespread than initially thought and its binding is associated with non-cancer cell lines.

With the enhanced mapping power afforded by long reads, we sought to determine if CTCF binding in mitosis occurs at loci inaccessible to short read technologies. Previous studies suggest that during mitosis in HeLa cells CTCF binds pericentromeric regions containing a shorter submotif found only in a subset of CTCF motifs known as M2 ^37^. We then searched for all M2 motifs in centromeric and pericentromeric DNA sequences and found that the overwhelming majority of M2 motifs occur in centromeric transition zones, which border the centromeres of chromosomes and frequently contain highly repetitive and difficult-to-map sequences ^39^. After averaging methyladenine around M2 sites across all subsets of centromeric regions we detected weak average CTCF binding at M2 sites specifically in centromeric transition regions and specifically during mitosis (Figures 2F, 2S1D-E).

A previous study of HeLa cells has also suggested that CTCF binding persists at the H19 locus during mitosis ^40^. We also sought to determine if haplotype-specific CTCF interactions can persist during mitosis. Although limited by the read depth of the experiment, we indeed observed that the H19 locus shows evidence of both CTCF binding on the maternal allele and increased CpG methylation on the paternal allele (Figure 2 S1F). However, further studies with higher depth datasets will be necessary to determine the full extent of haplotype-specific interactions during mitosis.

## Discussion

Here, we have developed, optimized, and validated DiMeLo-cito, a single-tube protocol for mapping protein-DNA interactions genome-wide. We show that DiMeLo-cito reproducibly maps CTCF sites with comparable or better performance compared to standard DiMeLo-seq (Figure 1B). We also demonstrate that DiMeLo-cito is compatible with ultra high molecular weight DNA extraction, for which we provide optimized protocols (Figure 1 S3B, Methods). Developing a one-tube protocol necessitated the replacement of the multivalent Protein A with monovalent anti-IgG nanobodies, which improves the efficiency of methyladenine deposition at target sites while reducing the radius of methylation (Figure 1D). The tighter labeling efficiency can be partially attributed to the smaller size of nanobodies (∼15 kDa) compared to Protein A (∼40 kDa), as the linker lengths between these domains in our Hia5 constructs are identical. Furthermore, because the rabbit nanobody tested here targets the Fab region of IgG, which directly contacts antigen, enzyme complexes may more closely align to the CTCF binding site when compared to Protein A, which instead binds at multiple sites on the antibody Fc region^21^.

A major motivation for developing DiMeLo-cito was to improve sample handling and recovery through the DiMeLo protocol. We achieved this by replacing wash steps with an affinity resin assembly known as an EMPaNADA. Enclosure of affinity resin into the EMPaNADA was important, as omission of this led to an ∼5-fold reduction in DNA recovery. Taking advantage of the enhanced efficiency of DiMeLo-cito, we were able to generate maps of CTCF binding from 100,000 cells, which is 10X lower than the lowest input used for DiMeLo-seq. Although we were able to generate data that suggests confident mapping of CTCF sites, we were limited in the depth achievable by ONT sequencing. As the performance and efficiency of ONT and other third-generation sequencing technologies improves we expect DiMeLo-cito to be positioned well to interrogate ever-smaller samples with greater depth.

The absence of wash steps also enables the interrogation of samples that are sensitive to centrifugation. Interrogation of mitotic chromatin binding is notoriously difficult due to the fact that mitotic chromosomes are highly vulnerable to fixation artifacts ^2^, necessitating the use of unfixed cells in mapping protocols. The absence of fixation, coupled with the fact that mitotic chromosomes are no longer enclosed in a nuclear envelope, renders chromosomes vulnerable to damage during repeated pelleting and wash steps. Using the wash-free DiMeLo-cito protocol, we unexpectedly found that CTCF binding during mitosis is widespread across chromosomes in GM24385 LCs (Figure 2). This observation requires the use of a wash-free protocol as standard DiMeLo-seq did not capture strong CTCF binding at these sites (Figure 2B). As a result, we speculate that centrifugation-based washes of mitotic chromosomes lead to greater chromosomal damage and/or displacement of CTCF binding that obscures its true binding distribution. This observation raises the possibility that the genomic binding of other important factors to mitotic chromosomes are also diminished by sample washing. Future studies that employ DiMeLo-cito and other methods that minimize sample manipulation will be necessary to clarify the true extent of mitotic chromatin binding. The use of affinity depletion within devices such as the EMPaNADA design introduced here offer a potentially generalizable approach to replace wash steps and enable sensitive studies in biochemical protocols. Indeed, we expect that the use of EMPaNADAs could readily be adapted to short-read mapping approaches such as CUT&RUN as a short-read alternative to DiMeLo-cito for the mapping of chromatin structure in sensitive samples.

How chromatin structure is inherited across cellular generations is a major outstanding question, as interphase chromatin structure is completely lost during the compaction and segregation of chromosomes during mitosis ^41^. Increasingly, studies suggest that a subset of critical chromatin regulators remain bound at their target sites during mitosis to “bookmark” these sites for rapid reconfiguration upon exit from mitosis ^42^. Although CTCF has been proposed to act like this in mouse studies, previous studies of CTCF binding in human cells have suggested that nearly all CTCF binding is lost during mitosis ^7^. However, many of these previous studies were done using transformed cancer cell lines. One study using single molecule imaging in fetal lung fibroblasts found that most CTCF molecules are unbound in mitosis, but this study did not measure the occupancy of CTCF sites on mitotic chromosomes^43^. By applying DiMeLo-cito to the non-cancerous GM24385 cell line, we unexpectedly found strong mitotic CTCF binding, which we validated by IF staining in unfixed GM24385 cells and H1 hESCs (Figure 2C). Consistent with previous studies, DiMeLo-cito showed that CTCF binding is largely lost during mitosis in HeLa cells. We validated this loss of binding by IF staining in HeLa, U2OS, and Ramos cells, which are all cancer-derived (Figure 2C). These observations raise the possibility that CTCF binding during mitosis is more widespread in human cells than initially suspected, and it is frequently disrupted in cancer cells.

Future studies employing DiMeLo-cito and other methods will be necessary to determine what mediates the apparent CTCF binding differences in various cell types. It has been demonstrated that phosphorylation of CTCF, which occurs during mitosis, reduces its affinity for targeted motifs ^44^, raising the possibility that the degree of CTCF phosphorylation may differ between cell types. Since patterns of CTCF binding instruct the folding of interphase chromosomes ^45^, binding of CTCF during mitosis may safeguard the building of interphase structures across cellular generations. This also raises the possibility that its abnormal loss during mitosis enables dysfunctional chromatin states that underlie disease. It is our hope that DiMeLo-cito will help to yield many additional discoveries in the field of chromatin biology.

## Materials and Methods

### Cell culture

GM24385 lymphoblastoid cells (Coriell Institute; mycoplasma tested) and Ramos B-Cell Lymphoma cells (ATCC; mycoplasma tested) were maintained in suspension using RPMI-1640 with l-glutamine (Gibco, 11875093) supplemented with 20% FBS (VWR 89510-186) and 1% penicillin–streptomycin (Gibco, 15070063) at 37 °C in 5% CO_2_. HeLa and U2OS cells were maintained in DMEM with l-glutamine (Gibco, 10564011) supplemented with 15% FBS (VWR 89510-186) and 1% penicillin– streptomycin (Gibco, 15070063) at 37 °C in 5% CO_2_. H1 cells were obtained directly from WiCell at passage 22. Cells were cultured according to WiCell’s protocol using Matrigel coating (Corning 354277), mTESR Plus (Stem Cell Technologies 100-0276) and Versene EDTA (Fisher Scientific A4239101). Briefly, cell culture surfaces were coated with Matrigel for 2 hrs or longer at 37 C in the tissue culture incubator. For passaging, cells were rinsed with Versene, incubated with Versene for 8.5 minutes at room temperature. Versene was aspirated, and then fresh mTESR Plus media was used to lift hESC colonies from the dish. Cells used for DiMeLo and DiMeLo-cito were washed twice with PBS by pelleting cells at 500xg for 5 minutes and aspiration of the supernatant before resuspension. After washing, cells were counted and aliquots into 1 million cell aliquots. Aliquots were spun down at 500xg for 5 minutes in 2mL DNA Lo Bind tubes (Eppendorf 022431048) before aspirating the supernatant and flash freezing in liquid nitrogen. Samples were stored at −80°C until use.

For synchronization of GM24385 LCs in M phase, we performed a double thymidine block followed by release into nocodazole as described previously ^46^. Briefly, GM24385 cells were incubated for 16 hours with media containing 4 mM thymidine (ThermoFisher, A11493.22) and then washed into pre-warmed media without thymidine. After 6-8 hour incubation, 4 mM thymidine was added to the media and cells were again incubated overnight. The following day, cells were washed into fresh pre-warmed media without thymidine and incubated for 6-8 hours before addition of 40 ng/mL Nocodazole (Abcam. ab120630) overnight. The following day, cells were twice washed into HBSS (Santa Cruz sc-391061A) containing 1 µM MG132 (Stem Cell Technologies 73262). Cells were then pelleted and supernatant aspirated before flash freezing in liquid nitrogen and storing at −80°C. For synchronization of U2OS and HeLa cells, cells were incubated overnight once with 4 mM Thymidine, washed into fresh media and incubated for 6-8 hours before addition of 40 ng/mL Nocodazole and incubation overnight. Mitotic cells were released from the plate by gentle shake off and collection of the media into HBSS containing 1 µM MG132.

### Immunofluorescence, imaging, and flow cytometry

For immunofluorescence staining of cell pellets, frozen samples were resuspended in 100 µL permeabilization buffer (20 mM HEPES pH 7.5 150mM NaGlutamate 0.5 mM Spermidine 0.1% w/v BSA 1X Roche Complete protease inhibitors 0.1%Tween 20 0.02% digitonin) and incubated for 5 minutes on ice. Primary antibody (see below for information) was added at 1:100 dilution and incubated for 2 hours at 4°C on a rotating mixer. Samples were washed with 3X 500 µL with TBST by pelleting at 300xg for 5 minutes. Samples were then incubated for 1 hour with Alexafluor 549 secondary antibody (Thermofisher A-11012) diluted in permeabilization buffer without digitonin and then washed 3X with 500 µL TBST. For imaging, cells were then resuspended in vectashield with DAPI (vector labs, H-1200-10) and pipetted onto a glass slide pre-coated in polylysine, covered with a glass coverslip, and sealed with nail polish prior to confocal imaging. For flow cytometry, samples were resuspended in HBSS+1% serum and 1 ng/mL DAPI, strained with a 40 µm strainer, and run on a Sony Sorter SH800S in analysis mode. Cell nuclei were gated based on FSC/BSC of lymphocytes. For h1 hESCs, cells were permeabilized, stained, and washed, according to the same buffers as above but while remaining adhered to matrigel coated confocal imaging plates (MaTEK). This was done by incubation with gentle rocking at 10rpm at 4°C so as not to release adhered cells and solutions were removed vacuum aspiration instead of pelleting. Samples were coated in vectashield with DAPI prior to confocal imaging.

ROI’s containing mitotic chromosomes were manually identified and segmented using imagej. For each ROI’s, a 7X7X3 pixel median filter was performed on the DAPI channel, followed by intensity thresholding and object size filtering to identify pixels belonging to the mitotic chromosomes of individual cells (all operations performed in 3D). The resulting 3D pixel mask was used to query the CTCF signal on the mitotic chromosomes. To estimate the cytoplasmic CTCF intensity, a 2×2×2 pixel mask shell was generated around the chromosome mask. For each mitotic cell, the mean CTCF signal was calculated for the mitotic chromosome mask and the cytoplasm shell. The ratio of these intensities is plotted in Figure 2C. A two-sample tailed t-test is calculated between the secondary control condition and each cell line.

### Expression and purification of Hia5 constructs

Protein A-Hia5 was expressed and purified as previously described^10^. MBP-Nanobody-Hia5 constructs were synthesized and cloned into rhamnose-induction expression vectors by ATUM. The nanobody TP896 targeting rabbit IgG Fab and TP1107 targeting mouse IgG Fc were used for anti-rabbit and anti-mouse Hia5 respectively. T7 Express lysY/Iq (NEB C3013I) cells were transformed with expression plasmid according to manufacturer specifications and plated on Kan-30 plates. Individual colonies were picked and grown overnight in autoclaved LB media (RPI) starter cultures overnight at 37°C. Starter cultures the next day were diluted 100X in autoclaved TB (RPI) and grown at 37°C with shaking at 200 rpm in 2.8 L baffled flasks to OD∼0.6-0.8. Cultures were then cooled to 18°C before addition of 0.1% rhamnose to induce expression. After incubation for 16-20 hours at 18°C cells were pelleted by centrifugation at 4000xg for 20 min at 4°C. Supernatant was aspirated and cell pellets were stored at −80°C until processing To purify MBP-Nanobody-HIa5, cell pellets from 3-6L of culture were resuspended in 100mL lysis buffer (20 mM Tris pH 8.0. 300 mM NaCl, 10% glycerol, 10 mM Imidazole, 1X Roche Complete Protease inhibitors, 1U/mL Benzonase) on ice and lysed using a C5 Emulsiflex microfluidizer (Avestin). Cell lysates were clarified by centrifugation at 16000xg for 30 minutes. Clarified lysate was applied to a 5 mL histrap FF column (Cytiva, 17525501) using a luer lock syringe fitted with an adapter. The histrap column was then washed with 5 additional column volumes of lysis buffer with 500 mM NaCl followed by an additional 5 column volumes of lysis buffer. Samples were then eluted with elution buffer (20 mM Tris pH 8.0. 300 mM NaCl, 10% glycerol, 500 mM Imidazole, 1X Roche Complete Protease inhibitors) and dialyzed into IEX Buffer A (20 mM Tris pH 8.0. 50 mM NaCl, 10% glycerol 2 mM DTT) overnight. The next day dialyzed protein was loaded on a hiTrap heparin column (Cytiva, 17040703) using an Akta Pure system (Cytiva). The column was washed with 5 column volumes IEX Buffer A before being subjected to a linear gradient from 0-50% Buffer B (Buffer A with 1M NaCl) over 20 column volumes with fraction collection. Fractions containing target protein were identified by UV Chromatography at 280 nm and confirmed by reducing SDS-PAGE. Fractions containing the target protein were pooled, concentrated using Amicon 30 kDA MWCO concentrators (Millipore) and injected onto a Superdex 200 10/300 increase column (Cytiva, 28990944) equilibrated in SEC Buffer (20 mM Tris pH 8.0. 300 mM NaCl, 10% glycerol, 2 mM BME). Major peak fractions were verified to contain highly pure protein by reducing SDS-PAGE. Protein was then concentrated to ∼1 mg/mL by absorbance at 280 using the sequence predicted extinction coefficient from expasy protparam assuming all residues are reduced. Protein was aliquoted and flash frozen in liquid nitrogen before storage at −80°C.

### Restriction enzyme methyltransferase assays

Methyltransferase assays were carried out on 30 ng of dsDNA oligos of 100bp containing a single Dam site at the center of the sequence. They were methylated in the presence of DiMeLo activation buffer (15 mM Tris pH 8.0 60mM KCl/acetate/glutamate, 15 mM NaCl/acetate/glutamate, 0.05 mM Spermidine, 1 mM EDTA, 0.5 mM EGTA, 0.1% BSA, 800 µM SAM). Reactions were run for 1 hour at 37°C before terminating the reaction by heat inactivation for 5 min at 85°C. DNA from reactions were purified using 2 reaction volumes of Ampure XP beads (Beckman Coulter, A63881) according to the manufacturer’s protocol. Purified oligo was then digested overnight with 20U DpnI or DpnII (NEB) in 1X cutsmart buffer overnight at 37°C. Digestion reactions were then resolved on a tapestation using the D1000 kit or using an 8% polyacrylamide 1XTBE gel. Gel samples were quantified in ImageJ by densitometry.

### Preparation of EMPaNADAs

Approximately 30×20mm rectangles of PVDF membrane (1620261 BioRad) were cut and wet in nuclease-free water. Strips of membrane were folded in half “hamburger style” without creasing and sealed together at the edges of the fold using a heat-sealing iron (Alitamei/ASIN B07TSN8Z6N). Another edge of the EMPaNADA was sealed to form a pocket. Approximately 120µL of Protein A/G (Cytiva) or anti-MBP (NEB) paramagnetic affinity resin slurry were equilibrated into PBS by magnetic precipitation of beads and resuspension in 150 µL buffer three times. Bead slurry was then pipetted into the pocket and sealed on the third edge to fully enclose the EMPaNADA. To verify the absence of leaks, EMPaNADAs were incubated in a 5 mL tube containing PBS for at least 2 hours and watched for no accumulation beads at the bottom of the tube.

### DiMeLo and DiMeLo-cito workflows

DiMeLo-seq was performed according to protocols.io version 2 (https://www.protocols.io/view/dimelo-seq-directed-methylation-with-long-read-seq-n2bvjxe4wlk5/v2). For DiMeLo performed with glutamate salts the activation buffer replaced NaCl and KCl with the same concentrations of Na Glutamate and K Glutamate respectively with otherwise identical compositions. DiMeLo-cito was initiated by addition of 50 µL Buffer 1 (20mM HEPES pH 7.5 150mM NaGlutamate 0.5 mM Spermidine 0.1% w/v BSA 1X Roche Complete protease inhibitors 0.02% digitonin) to frozen cell pellets on ice to permeabilize cells. Samples were allowed to incubate for 5 min on ice and resuspended by gentle titration with a 200 µL pipette with a wide bore pipette tip before addition of 1µg of antibody and incubation with end over end rotation on a Hula mixer at 10rpm for 2 hours. Afterwards 50µL of Buffer 2 (20 mM HEPES pH 7.5 150 mM NaGlutamate 0.5 mM Spermidine 0.1% w/v BSA 1X Roche complete protease inhibitors 0.1% tween 20) was added along with 3:1 molar ratio MBP-nanobody-Hia5 ( ∼200 nM in 100µL assuming antibody molecular weight is 150 kDa). Samples were placed on a rotating hula mixer and incubated for 1 hour at 4°C at 10rpm. After incubation, 900 µL Buffer 3 (15 mM Tris pH 8.0 60 mM Kglutamate, 15 mM Naglutamate, 0.05 mM Spermidine, 1mM EDTA, 0.5 mM EGTA, 0.1% BSA) was added to the sample along with a pre-assembled EMPaNADA and incubated at room temperature on a nutating mixer for at least 2 hours. Reactions were regularly inspected to ensure that the EMPaNADA was immersed fully in the reaction buffer during incubation. Activation of the reaction was carried out by addition of 2.5 µL of 32 mM S-AdenosylMethionine (NEB) and 1 µL of 1 M DTT and incubation was carried out in the presence of the original EMPaNADA for 2 hours on a nutator at 37°C. After one hour of incubation reactions were supplemented with an additional 2.5 µL of 32 mM SAM. After 2 hours total of incubation, EMPaNADAs were removed and reactions were stopped by the addition of 2U Proteinase K (NEB), 1U RNase A (NEB) and 10 µL 10% SDS (Thermo). Samples were then incubated for 5 min at 65°C to digest protein and RNA before being subjected to DNA extraction. Antibodies used in this study were Rabbit anti-CTCF (Abcam) Mouse anti-CTCF (Thermo 1D11) Rabbit anti-H3s10ph (CST #53348) Rabbit Igg control(ab37415) and Mouse IgG Control (ab37355).

### DNA Extraction and Agilent Femto Pulse Analysis

DNA extraction was performed using the NEB monarch genomic and UHMW DNA extraction kits with minor modifications. To purify normal molecular weight DiMeLo-cito samples with 1 million cells or greater were applied to genomic spin columns after addition of 800 µL of kit binding buffer and applied to a single spin column in two separate spins. Samples were then processed according to the manufacturer’s specifications. For ultra high molecular weight purification, samples of ≥1 million cells were treated with 400 µL binding buffer and 600 µL to precipitate DNA onto beads. For samples with < 1 million cells as input, DiMeLo-cito samples were concentrated by SpeedVac set to 50°C and using infrared light to ∼200 µL before proceeding with the manufacturer’s protocol for all steps after the lysis step.

Ultra high molecular weight DNA was also extracted from DiMeLo-cito samples according to an in-house protocol. Samples containing ≥1 million cell inputs were directly taken into purification while smaller cell inputs were first concentrated by speedvac to ≤200 µL. Borosilicate glass beads of 3mm diameter (Chemglass, CG110102) were then added to the samples with 4 being added to those with larger cell inputs and 2 being added to those with smaller inputs. Samples were gently inverted with 0.4 volumes 5X Binding buffer (100mM Bis Tris pH 7.0 1% Domiphen Bromide, 10% PEG8000) and 600µL isopropanol. Samples were incubated for 10 min rotating at 30 rpm on a rotating mixer with samples on their side. Samples were visually inspected for the binding of DNA to large silica beads with a glassy gel of DNA fully enveloping the beads. Solution was carefully aspirated away from the beads taking care not to aspirate away viscous DNA. Samples were then washed twice each with gentle inversion after addition of 500µL using three wash buffers: Wash 1 (20mM BisTris pH 7.0 50%v/v 1,3 butanediol 2M Guanadine HCL, 2%w/v PEG 8000), Wash 2 (20mM BisTris pH 7.0 50% 1,3 butanediol, 2%w/v PEG 8000), and Wash 3 (20mM BisTris pH 7.0 50mM NaCl, 5%w/v PEG 8000) buffers. After the final wash with Wash 3, beads were carefully transferred to an empty bead retaining spin column (NEB, T3004L) placed in a fresh 2mL DNA lo bind collection tube and lightly pulse spun in a bench top centrifuge to remove residual wash buffer. Beads were then transferred to an empty 2mL DNA lo bind tube and DNA was eluted with 100 µL 10 mM Tris pH 9.0 1 mM EDTA 0.5 M 6-aminocaproic acid by incubation at 37°C for 10 minutes. The resulting eluate is highly viscous and is then separated from beads by transferring bead and sample mixture into a fresh, empty DNA lo bind tube. To quantify ultra high molecular weight DNA, 10 µL aliquots were carefully pipetted so as not to shear the stock DNA and placed into a fresh 1.5 mL DNA lo bind tube along with a single 3 mm diameter silica bead. The sample was then pulse vortexed prior to measurement to obtain low viscosity samples that can be accurately quantified on a nanodrop or qubit system. Samples for Femto Pulse analysis were prepared according to the 165 kb genomic DNA analysis kit.

### Oxford Nanopore Sequencing and Analysis

DNA sequencing libraries were prepared using the rapid barcoding v14 (RBK114-96) and native barcoding v14 (NBD114-96) kits according to manufacturer specifications. Sequencing libraries were sequenced on Promethion R10.4 flow cells and inline basecalled with dorado v0.7 on Nvidia A100s using 6mA and 5mCpG super accurate modified basecalling models. BAM files produced from basecalling were merged and aligned to either the CHM13v2.0T2T genome or the Hg002 haplotype resolved genome using minimap2 with the ont preset and then sorted and indexed using samtools. Further analysis was carried out using the dimelo v2 python package (https://github.com/streetslab/dimelo_v2) and plotted in matplotlib. Unless otherwise stated all pileup and enrichment plots generated by the dimelo package were using and Ml tag threshold score of 225 (0.88 confidence) while all single read plots were generated using a threshold score of 250 (0.98 confidence). BED files of CTCF peaks were generated from GM12878 and HeLa Encode datasets. Bed files for CTCF M2 motifs were found using fimo with default parameters and intersected with censat annotations using bedtools.

## Data & Code Availability

Source data and code will be made available upon publication or request to the corresponding author. Nb-Hia5 plasmids will be deposited in Addgene and are currently available upon request to the corresponding author.

## Author Contributions

N.G. and N.A. designed the study. N.G. performed and analyzed all experiments included in the manuscript. N.G. C.C. K.S. J.P.S. generated and purified Hia5 constructs. N.A., S.L., and D.R. performed initial restriction assays used to design DiMeLo-cito. O.D.L., J.M., and A.M. developed the dimelo analysis package and assisted in analysis. J.M. and A.M. performed initial experiments involving reducing agents. J.M.F. assisted in collection and analysis of imaging data. A.S., A.F.S., and N.A. supervised the research. N.G. and N.A. wrote and edited the manuscript.

## Acknowledgements

We would like to thank the Stanford Protein and Nucleic Acids facility in the Stanford School of Medicine for helping operate and maintaining the Agilent FemtoPulse system. We would also like to thank all members of the Altemose laboratory for helpful discussions and comments in preparation of this manuscript.

## Supplemental Figure

**Figure 1 S1.**
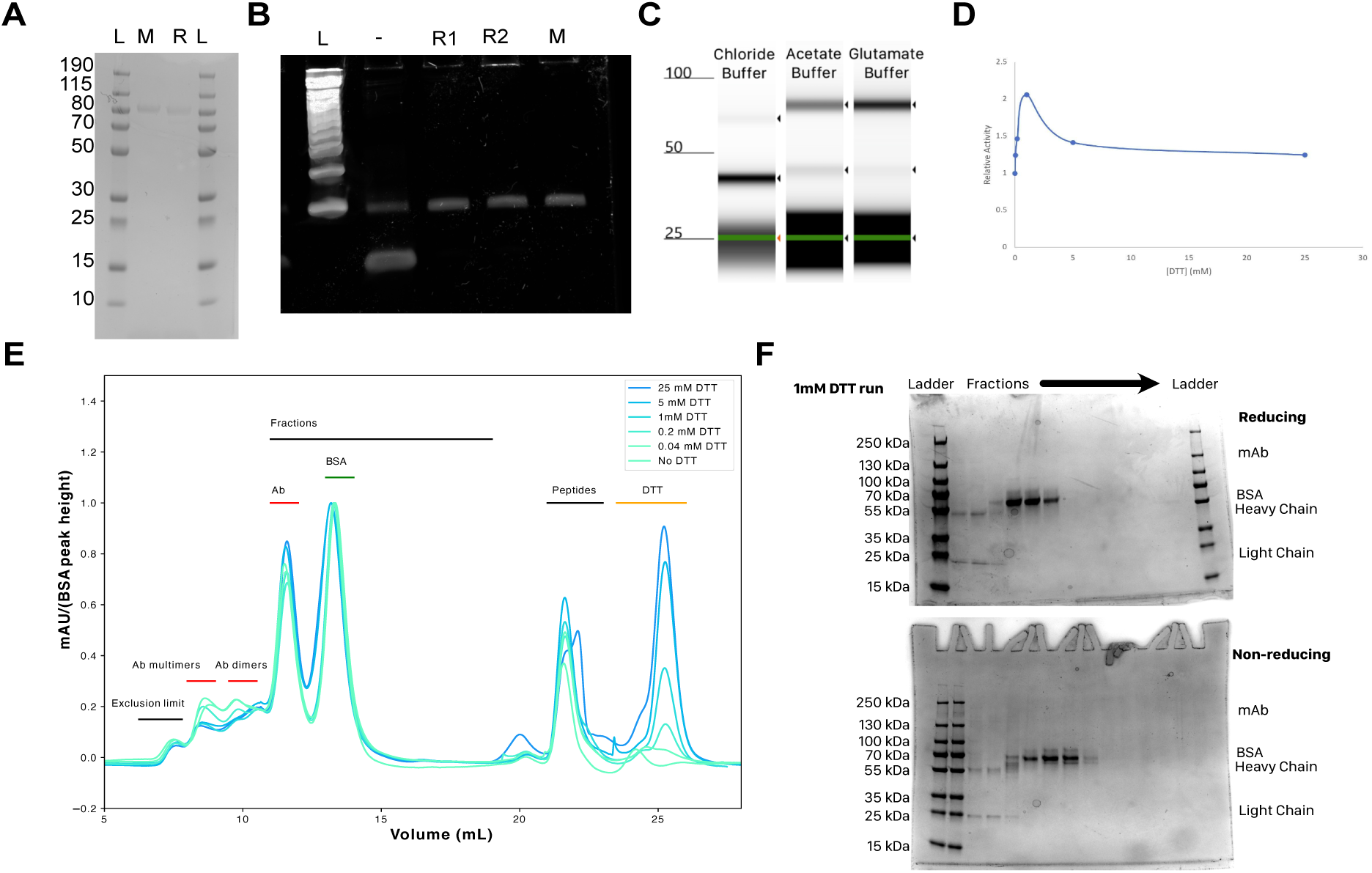
Optimization of Hia5 reaction conditions. **A**. SDS-PAGE of purified MBP-anti-mouse-Hia5 (M) and MBP-anti-rabbit-Hia5 (R). **B.** Polyacrylamide gel of a restriction protection methyltransferase assay with DpnII for MBP-anti-mouse-Hia5 (M) and two preparations of MBP-anti-rabbit-Hia5 (R1/R2) in glutamate buffer. Uncut fragments indicate higher degrees of methylation. **C.** Electropherogram from a Tapestation D1000 showing a restriction protection assay using MBP-anti-rabbit-Hia5 in DiMeLo activation buffer with the indicated counterion. **D.** Relative protection of DNA in a restriction protection assay using chloride buffer with increasing concentrations of DTT. **E.** Chromatogram at A280 from injections of anti-CTCF antibody onto an S200 increase 10/300 size exclusion column after incubation at 37C for 1 hour with increasing concentrations of DTT. Locations of the eluting species are marked above peaks. **F.** Reducing and Non-reducing SDS-PAGE of fractions eluting from size exclusion chromatography of Antibody incubated with 1mM DTT prior to injection.

**Figure 1 S2.**
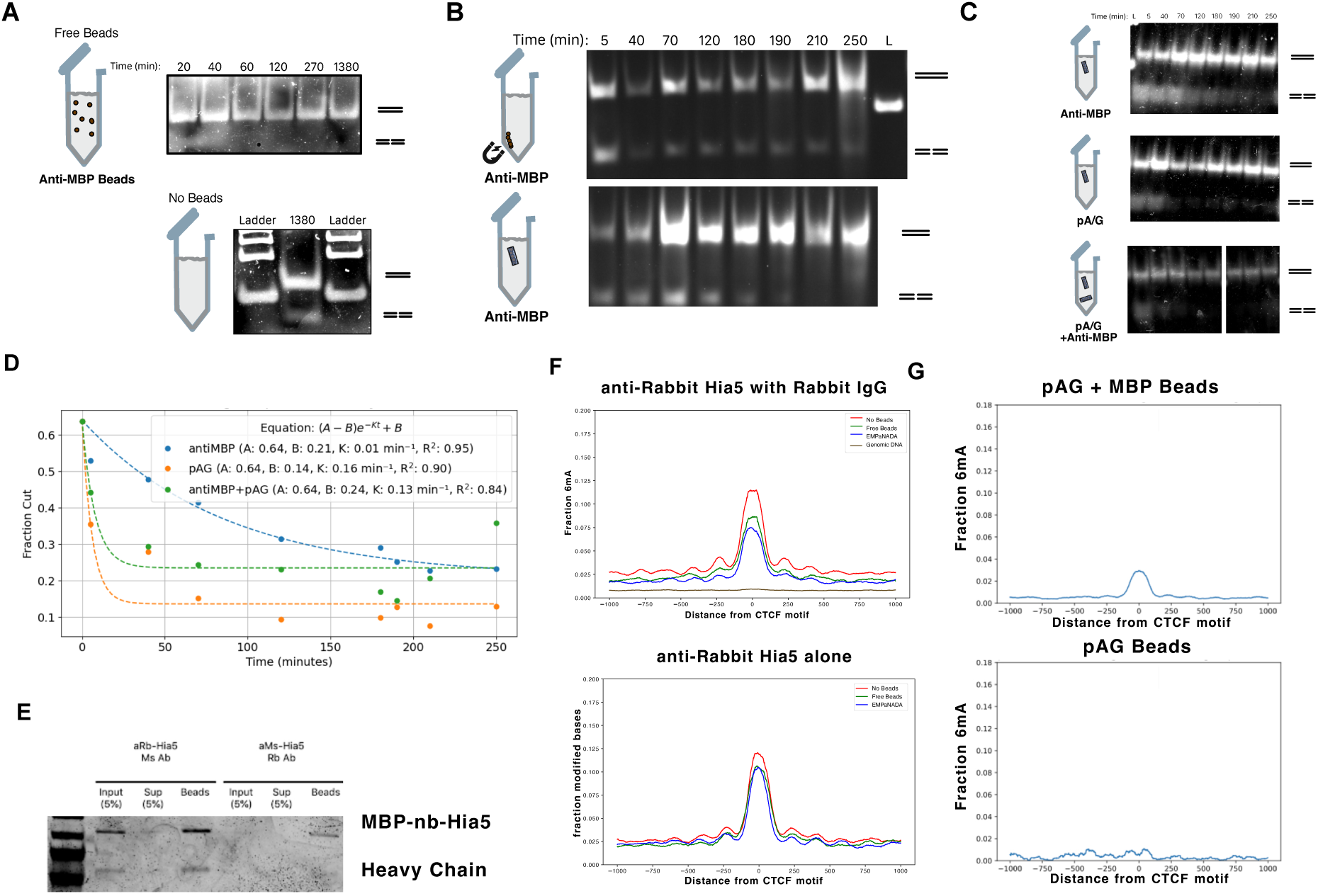
Optimization of free-enzyme depletion. **A.** Kinetics of antibody enzyme complex depletion with free beads. Anti-MBP beads were incubated with precomplexed antibody-Hia5 complexes (1:3) at room temperature for 5 minutes. Supernatants retrieved at the indicated time were then subjected to a restriction enzyme methyltransferase assay using DpnI where higher amounts of cutting indicate more methyltransferase activity. **B.** Kinetics of antibody enzyme complex depletion by restriction enzyme methyltransferase assay with DpnI comparing incubation with paramagnetic anti-MBP beads enclosed in an EMPaNADA or retained against the wall of the tube with a magnet using a DpnI. **C.** Kinetics of antibody enzyme complex depletion by restriction enzyme methyltransferase assay comparing incubation with a single Anti-MBP EMPaNADA, a single pA/G EMPaNADA, or both a single anti-MBP and pA/G EMPaNADA. **D.** Fits to a single exponential decay for the depletion reactions in C. **E.** Precipitation of the indicated nanobody-Hia5 construct onto pA/G beads. Samples were also incubated with a nonbinding antibody as a positive control. **F.** DiMeLo-cito in GM24385 LCs with pA/G bead EMPaNADAs using anti-rabbit-Hia5 in the presence or absence of control antibody sequenced to ∼1X coverage. Profile plots centered at CTCF peaks were generated with a 50 bp smoothing window. G. DiMeLo-cito using anti-rabbit-Hia5 with control antibody and either a single pA/G EMPaNADA or a pA/G and anti-MBP EMPaNADA. Samples were sequenced to ∼0.1X coverage and profile plots centered at CTCF peaks were generated with a 50 bp smoothing window.

**Figure 1 S3.**
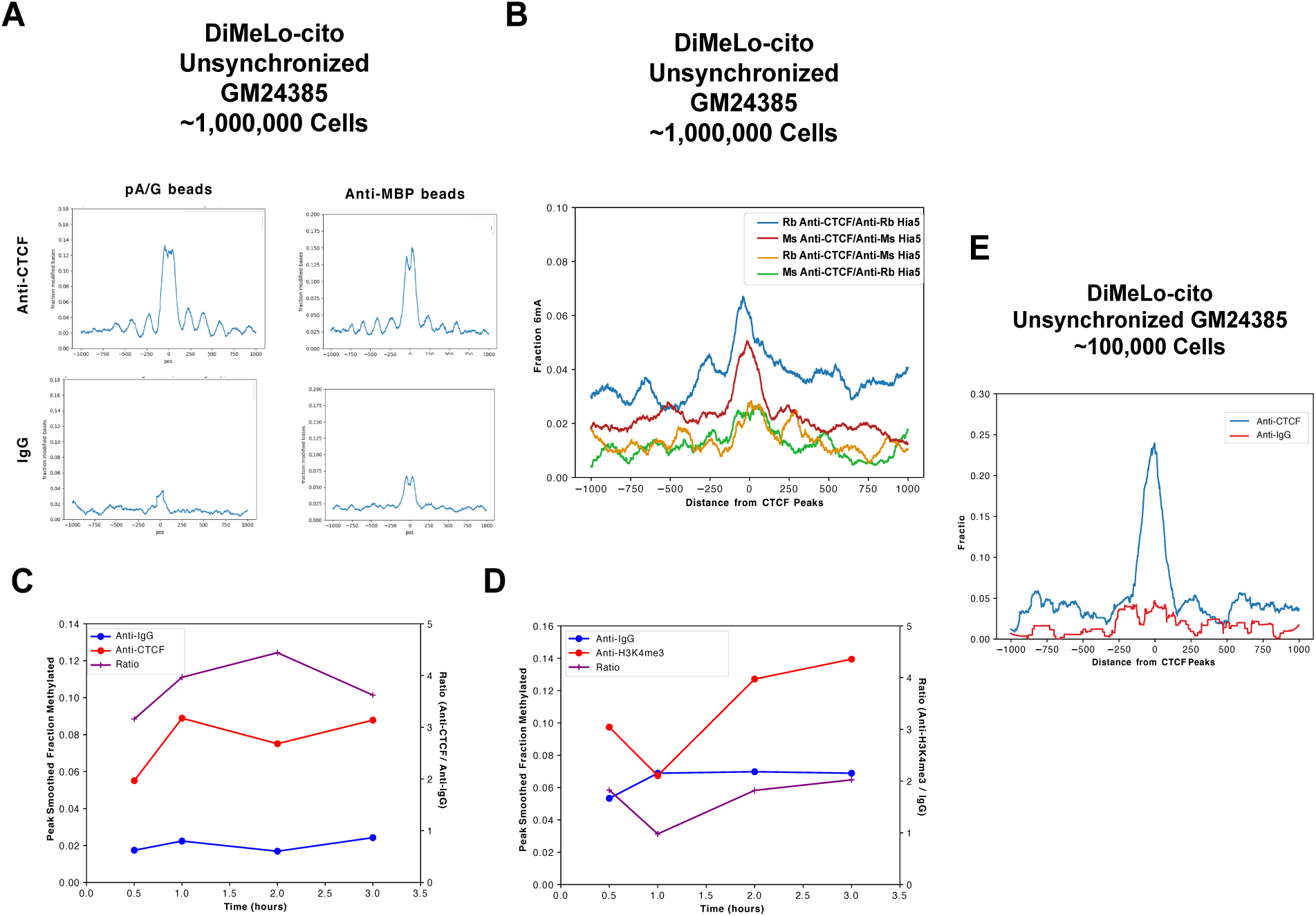
Optimization of DiMeLo-cito. **A.** Comparison of DiMeLo-cito using CTCF targeting or non-targeting with 1 million cells using pA/G or anti-MBP paramagnetic beads sequenced to ∼0.1X coverage. **B.** Comparison of DiMeLo-cito using rabbit or mouse CTCF targeting antibody combined with cognate or noncognate antibody. Samples were sequenced to ∼0.1X coverage and profile plots centered at CTCF peaks were generated with a 50bp smoothing window. **C.** Kinetics of methyladenine deposition by anti-rabbit-Hia5 during DiMeLo-cito at CTCF sites using CTCF-targeting or non-targeting antibody. Each point is the peak intensity from profile plots centered at CTCF sites from a single DiMeLo-cito reaction incubated for the indicated time during the activation step. Each reaction was sequenced to ∼0.1X coverage. **D.** Kinetics of methyladenine deposition by anti-rabbit-Hia5 during DiMeLo-cito at the top quartile of transcription start sites by RNA-seq using H3K4me3-targeting or non-targeting antibody. Each point is the peak intensity from 50 bp smoothed profile plots centered at H3K4me3 sites from a single DiMeLo-cito reaction incubated for the indicated time during the activation step. Each reaction was sequenced to ∼0.1X coverage. **E.** DiMeLo-cito targeting CTCF or a nontargeting control performed with 100,000 cells. Plot is smoothed with a 50 bp sliding window.

**Figure 1 S4.**
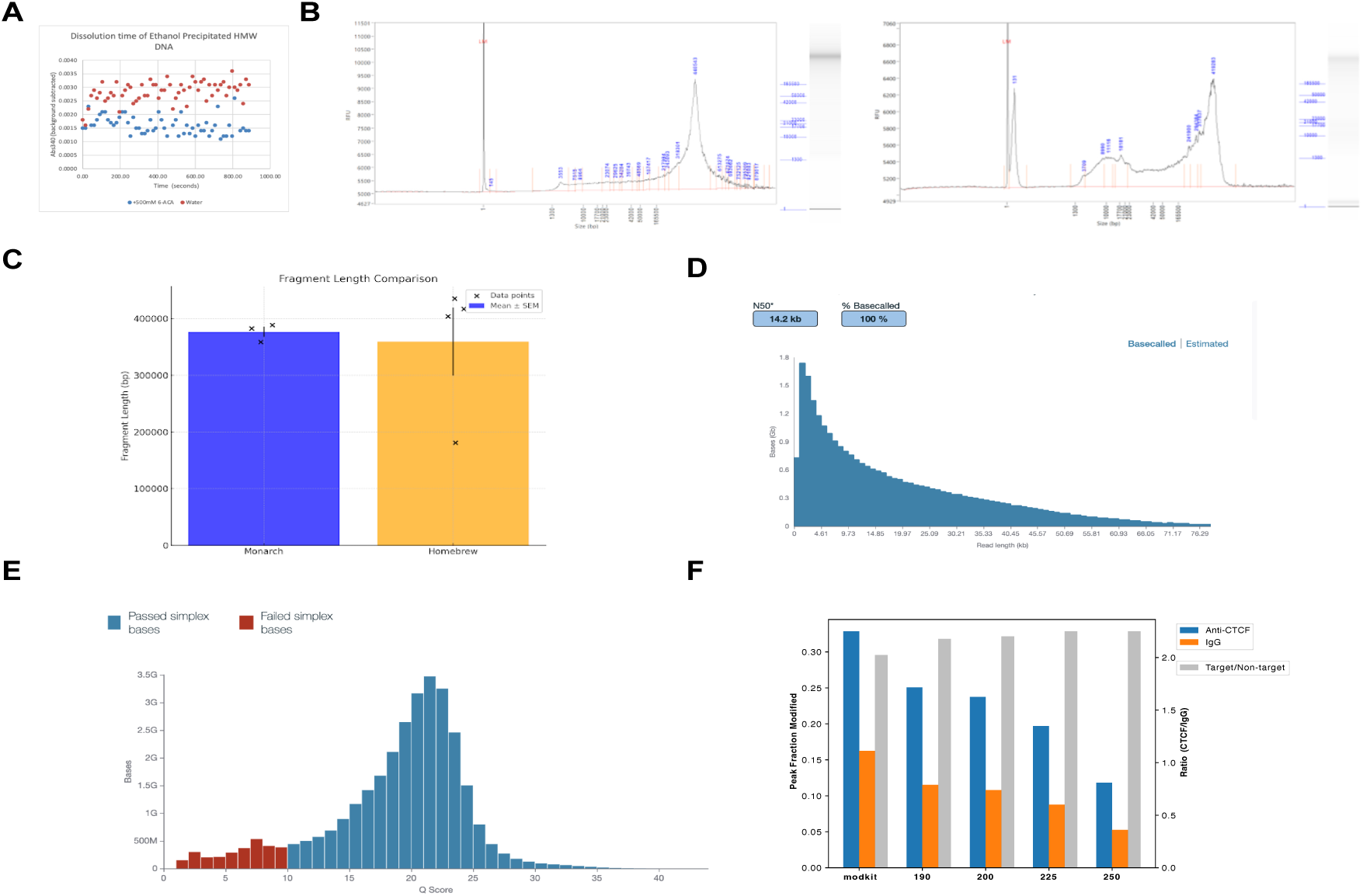
Properties of DiMeLo-cito DNA extraction and sequencing. **A.** Dissolution kinetics of precipitated ultra high molecular weight genomic DNA extracted from HeLa cells monitored by light scattering at 340 nm. The time course compares incubation at room temperature with nuclease free water or with 0.5M 6-ACA pH 9.0. **B.** Screenshots from Femto Pulse runs of DiMeLo-cito samples extracted using NEBs Monarch HMW genomic DNA kit or using an in-house method (see methods). **C.** Median fragment length distributions from samples extracted using the NEB kit or in-house (homebrew) method. **D.** Fragment length distributions from library preparation by ligation sequencing of HMW DNA extracted using the in-house method and sheared to generate smaller fragments. Sequencing data from this run was used in the generation of the 30X coverage dataset presented in Figure 1**. E.** PHRED Q-score distributions of basecalls using the super accurate modified base calling models from the same run as D. **F.** Plots of methyladenine enrichment from DiMeLo-cito at CTCF sites using CTCF-targeting or nontargeting antibody and the ratio of the two at various modification confidence thresholds. The x axis values correspond to the chosen minimal Ml cutoff score to be considered methyladenine or a cutoff score automatically chosen from the data by modkit within the dimelo package.

**Figure 2 S1.**
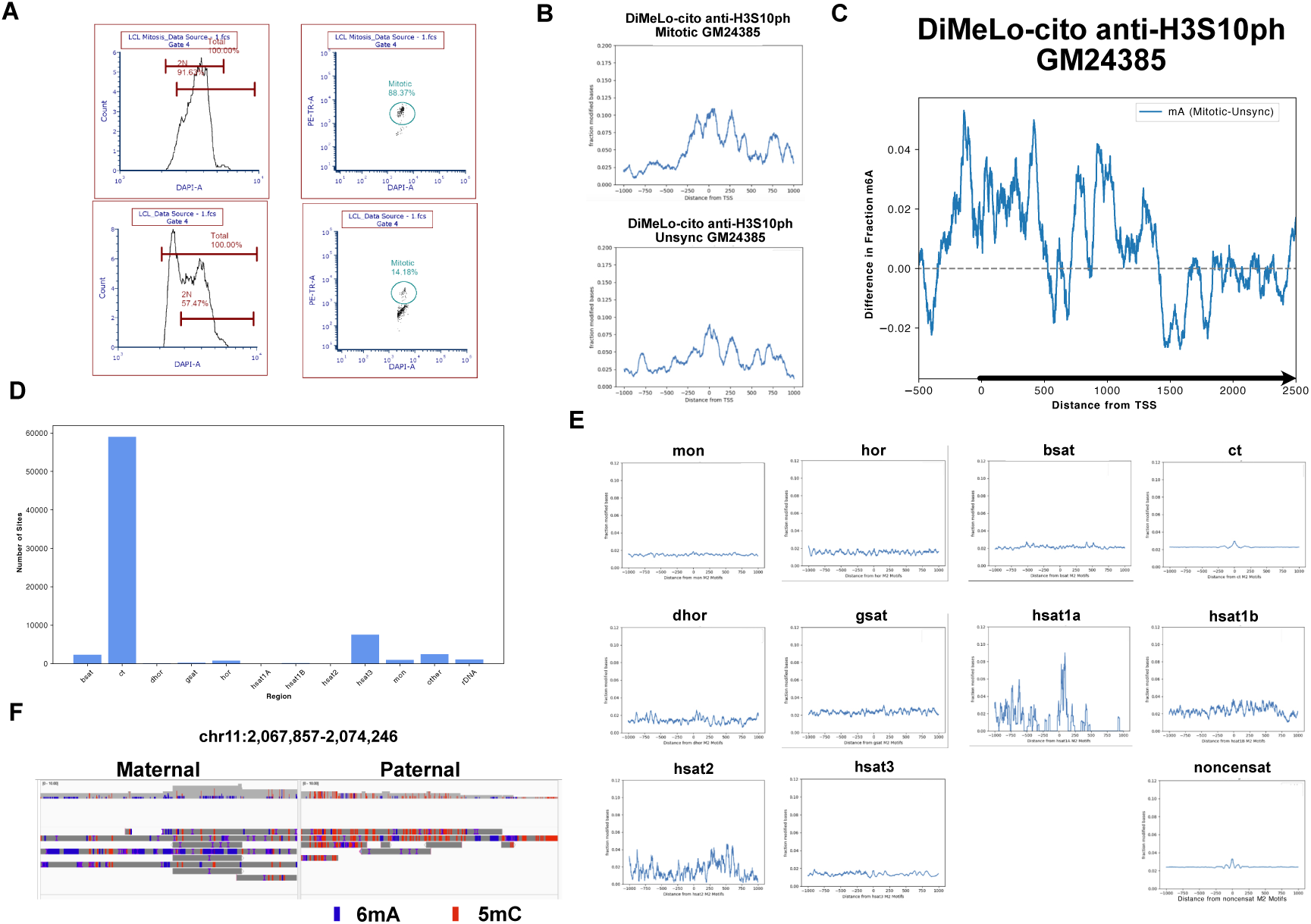
Validation of mitotic enrichment and distributions of centromeric CTCF in GM24385 LCs by DiMeLo-cito. **A.** Flow cytometry quantifying the fraction of cells past S phase by DAPI staining (left) and the fraction of cells actively in mitosis (right) by IF staining of H3S10ph. B. Enrichment profile plots of methyladenine at active transcription sites using H3s10ph targeted DiMeLo-cito in either mitotic (top) or unsynchronized GM24385 LCs (bottom) both smoothed with a 50 bp sliding window. **C.** Difference plot of profiles in B. **D.** Instances of CTCF M2 motifs identified by fimo in different centromere/satellite (censat) DNA annotated regions in the CHM13v2.0-T2T genome. **E.** Profile plots with 50bp smoothing of methyladenine at M2 motifs separated by censat annotations as well as all noncensat sites in the CHM13v2.0-T2T genome. **F.** Screenshot from IGV read browser at the H19 imprinting control region of anti-CTCF DiMeLo-cito in mitotic GM24385 LCs aligned to the HG002v1.1 genome.

**Figure 2 S2.**
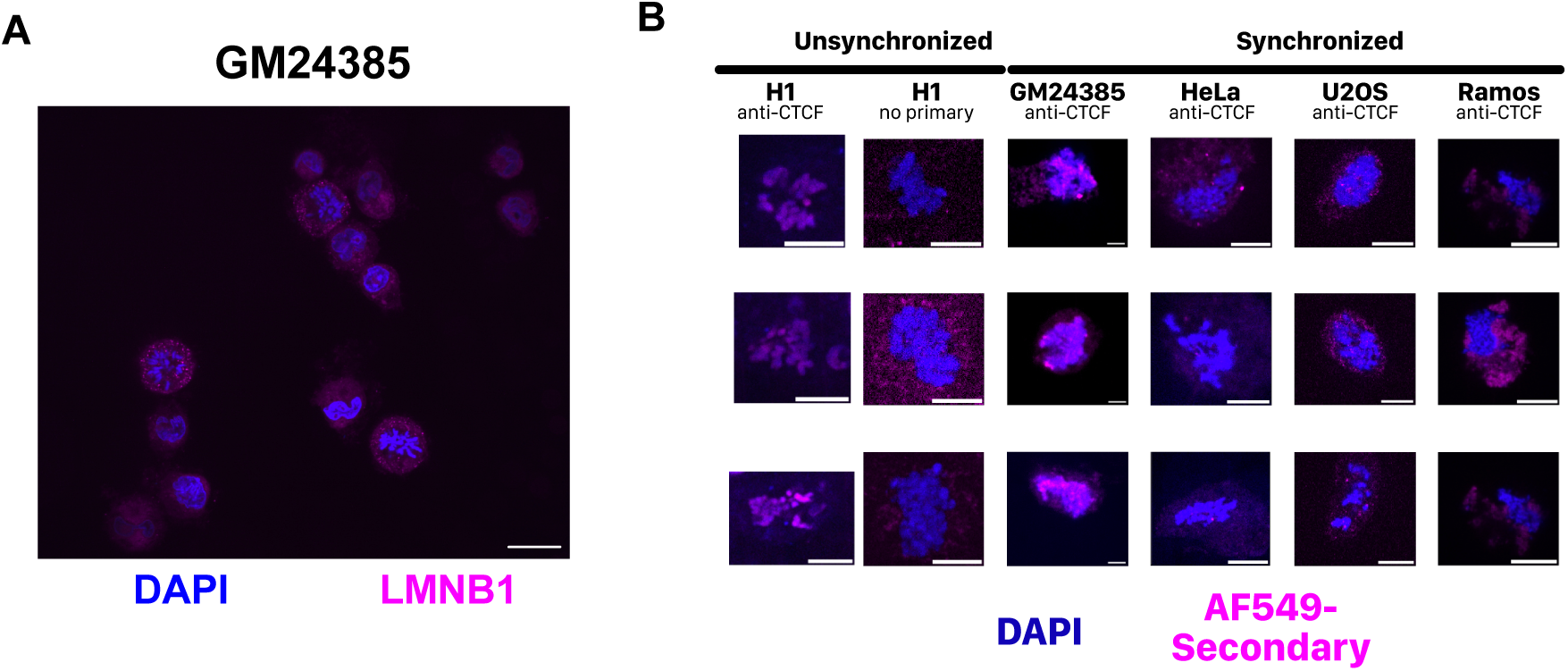
Variation in mitotic chromosome enrichment of anti-CTCF staining. **A.** Confocal imaging of mitotic synchronized GM24385 stained for the lamina. Scale bar corresponds to 15 µm. **B.** Confocal imaging of CTCF-stained or unstained IF samples in various unfixed cell lines chemically synchronized in mitosis or unsynchronized. Scale bar corresponds to 15 µm.

